# Computational aberration-corrected volumetric imaging of single retinal cells in the living eye

**DOI:** 10.64898/2026.03.21.712744

**Authors:** Guanping Feng, Dayn R Godinez, Zhongqiang Li, Stephanie Nolan, Hongkwan Cho, Elizabeth Kimball, Elia J. Duh, Thomas V Johnson, Ji Yi

## Abstract

The eye offers a unique non-invasive window for accessing single-cell level structures and functions of the central nervous system (CNS) throughout the retina. However, strong and space-varying ocular aberrations, along with limited volume rates, challenge large-scale cellular imaging in living eyes and stymie the full potential of possible biological and pathological studies in retina. Here, we present plenoptic illumination scanning laser ophthalmoscopy (PI-SLO), a 3D fluorescent retinal imaging modality that enables high-speed, widefield, volumetric single-cell imaging with low phototoxicity. By capturing multiple angular images of fluorescence signals from the entire volume, PI-SLO enables digital aberration correction and 3D imaging across a >20º FOV with >23 Hz volume rate. We leverage this structural and functional imaging modality to investigate three key aspects of CNS physiology through the living mouse retina, including: microglial process dynamics, vascular perfusion, and light evoked calcium fluxes in inner retinal neurons. PI-SLO is a versatile non-invasive platform for *in vivo* investigation of retinal and CNS physiology at the cellular level.

## 1. INTRODUCTION

The retina is a specialized neural tissue that sense light, processes visual signals, and enables visual sensation. Using the ocular optics as a lens, the retina provides the only optical window for direct *in vivo* observation of central nervous system (CNS) at a single cell level^1–6^, without invasive procedures such as craniotomy or tissue excision. The retinal circuitry is organized in highly stratified layers with diverse cell types connecting within a thin (200-300µm) three-dimensional (3D) volume^7^. This makes it an excellent target to study interactions across the CNS cell types without necessitating penetration through highly scattering deep brain tissue. Combined with widely available endogenous and exogenous fluorescent reporters and genetic editing toolbox, the retina enables molecular imaging with high specificity and functional readouts *in vivo* to allow broad range of biological studies, such as immune surveillance and responsivity^8–10^, neurovascular coupling^11^, and neural electrophysiologic activity under different stimuli^6,12,13^ just to name a few. Therefore, single-cell level volumetric retinal imaging is not only instrumental for studying retinal pathologies in preclinical models, but also for investigating fundamental mechanisms in basic neuroscience. Analogous to epifluorescence and confocal microscopy, fundus photography^14^ and scanning light ophthalmoscopy (SLO)^15,16^ have been widely used for *in vivo* retinal fluorescence imaging. However, neither modality provides sufficient spatial resolution and depth sectioning to resolve individual retinal cells in 3D, hindering the full potential of *in vivo* retinal imaging.

The physical barrier to achieve high-resolution *in vivo* imaging in the retina is refractive aberrations in the ocular optics, which degrades image lateral/axial resolution, image contrast, and signal to noise (SNR). To provide a context, healthy ocular optics typically exhibit significant wavefront errors, with root-mean-square (RMS) aberrations exceeding ∼0.3 µm in human^17,18^ and >1 µm in mice^19^, whereas diffraction-limited imaging typically requires <λ/14 RMS in microscopy (λ-wavelength). Importantly, ocular aberrations are eye-dependent, random, and unpredictable, preventing correction through optical design alone as in conventional microscope objectives. Originally developed for astronomy and later adapted for ophthalmic imaging, adaptive optics (AO) actively measures and corrects for each eye’s wavefront aberrations in real time, enabling near diffraction limited resolution^20–22^. It has been incorporated into nearly all optical ophthalmic imaging modalities to visualize cellular and even subcellular structures in both human and animal eyes^11,20,23–25^. Among AO ophthalmic imaging modalities, AOSLO is a powerful platform to study single cell level biology in the living retina by providing structural and functional fluorescent imaging of diverse retinal cell types, including immune cells^9,26,27^, retinal neurons ^6,11,25,28^, retinal pigmented epithelial cells^29–31^, pericytes^32^, and blood flow ^26,33^. Despite its resolution, monitoring large-scale retinal single cell dynamics in 3D remains challenging. First, the AO-corrected field of view (FOV) is limited by the isoplanatic patch size (∼1° or ∼300µm in humans ^34^, and ∼5° or ∼170µm in mice ^19^) which largely restricts imaging throughput; for example, the dendritic field of a retinal ganglion cell can span >160 µm in the mouse eye ^35,36^. Second, the confocal detection of AOSLO discards out-of-focus photons, requiring long integration times to resolve weak structures (e.g., >10 s to image microglia processes under 100 µW^9,37^). Moreover, sequential refocusing or axial scanning for volumetric imaging^11,19,25^ further increases phototoxicity, introduces temporal offset confounds between layers, and complicates motion registration.

To address this fundamental barrier of aberration, we use computational optics by engineering optical hardware to encode imaging depth and aberrations, then digitally reconstruct volume with aberrations corrected beyond the isoplanatic patch limit without hardware AO and sequential z-stacking. Our approach drew inspiration from light field microscopy (LFM), which has been successfully implemented for fast volumetric fluorescent imaging with high photon efficiency ^38–41^ and capability to infer and correct for optical aberrations ^42–44^. LFM collects photons from the entire illuminated samples while sampling both the spatial information and angular propagation of light rays from each voxels, known as the plenoptic function^38,39,45^. The plenoptic function encodes both the trajectories and origin positions of captured photons in 3D space, enabling joint reconstruction of 3D structure and pupil aberrations. Translating LFM to *in vivo* retinal imaging, however, has been unattainable due to multiple scattering across retinal layers, large and spatially varying ocular aberrations, weak signal with restricted light budget, and the impossibility of pre-calibration.

Here, we present a plenoptic illumination scanning light ophthalmoscope (PI-SLO) to capture plenoptic information of the eye, encoding both the depth of the retinal volume and the refractive properties of the ocular optics. Instead of using a microlens array (MLA) and camera in LFM^39,40,42^, PI-SLO captures plenoptic images by sequentially illuminating and scanning the retina from multiple incident angles with pupil modulation. The combination of selective angular illumination and point-scanning imaging improves robustness to multiple scattering in retinal tissue while enabling detection of weak fluorescence signals using sensitive photomultiplier tube (PMT). To pre-calibrate the highly aberrated ocular optics, a synthetic Hartmann-Shack wavefront sensor (Syn-HSWS) is deployed to measure the pupil aberrations and provide an accurate forward model prior. Together, PI-SLO enables digital correction of space-varying aberrations to achieve 3D single-cell imaging over a ∼700 × 500 × 160 µm volume in the living mouse eye, >12× larger than isoplanatic patch, at a >23 Hz volume rate. Photon efficiency is maximized by collecting emitted fluorescence from the entire volume without confocality and sequential z-stacking, reducing phototoxic exposure. Leveraging this technology, we demonstrated the capability of PI-SLO by imaging structures and functions of three key aspects of retinal and CNS physiology in living mouse eyes: 1) the transient dynamics of ∼100 single resident immune cells; 2) 3D retinal vasculature network across the entire inner retina; and 3) 3D signal neuron light-evoked intracellular calcium fluxes in GCaMP-expressing mouse retina. Our demonstrations establish PI-SLO as a powerful platform for large-scale *in vivo* investigation of retinal and CNS structure and function at single cell level.

## RESULTS

### Principle of PI-SLO for calibration-free computational 3D reconstruction with digital aberration correction

Unlike confocal AOSLO using full pupil aperture (Fig. 1a-i), PI-SLO deploys a configurable illumination aperture with a digital micromirror device (DMD) to sequentially scan 2D images through a translating sub-pupil aperture across the full pupil of the eye (Fig. S1). Illumination through an offsetting sub-pupil reduces the effective numerical aperture (NA) and generates a slanted and elongated point spread function (PSF) in the retina (Fig. 1b). Fluorescent photons emitted from the entire volume are captured by a photomultiplier tube (PMT) without a pinhole, enabling raster scanning across the retina to acquire angular parallax views of the 3D retinal volume (Fig. 1a-ii). The non-confocal configuration maximizes the photon collection efficiency compared to confocal AOSLO while leveraging the high-gain PMT to provide better sensitivity than a pixelated camera typically used in LFM. A lower excitation power can thus be used, which is particularly important for timelapse *in vivo* imaging where light budget is restricted.

**Figure 1.**
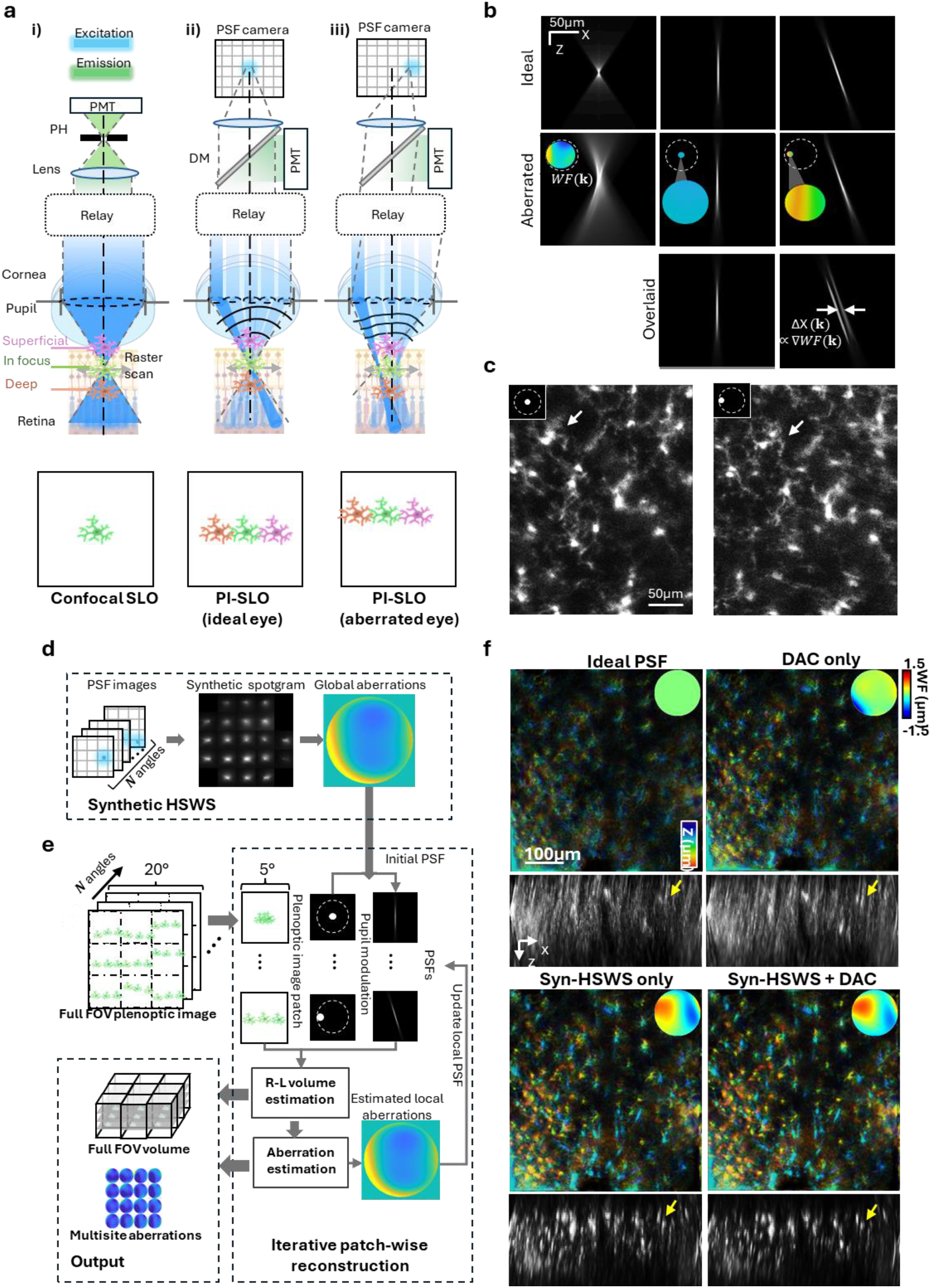
Principle of plenoptic illumination SLO. **a)** Schematic illustrations of different SLO illumination configurations (top) and their corresponding simulated images (bottom). **a-i**) Confocal SLO with full-pupil illumination. **a-ii-iii**) Plenoptic SLO with a single oblique sub-aperture illumination in an ideal eye **(a-ii)** and an aberrated eye **(a-iii**). **b)** Simulated three-dimensional point spread functions (PSFs) for different illumination configurations: full pupil (left column), central plenoptic angle (middle column), and off-axis plenoptic angle (right column). Unaberrated (top row) and aberrated (middle row) PSFs are shown for direct comparison. Aberration-induced lateral deviations of the plenoptic PSFs are highlighted by overlaying aberrated and unaberrated profiles (bottom row). **c)** *In vivo* plenoptic SLO images of GFP-expressing microglia acquired under two different illumination angles in a living *Cx3cr1-GFP* mouse. Arrows highlighted the lateral shifts of the same cells across angles revealing multiplexed depth and wavefront information. **d)** Illustration of synthetic Hartmann-Shack wavefront sensing. **e)** Illustration of the computational reconstruction routine. **f)** Comparison of *in vivo* 3D image reconstructed under different forward modes: ideal PSFs without aberration correction, only using digital aberration correction (DAC), PSFs generated by syn-HSWS, and combined syn-HSWS with DAC. The image was captured in a living *Cx3cr1-GFP* mouse eye in the laser-induced CNV model 2 weeks after laser injury. In each panel, the max intensity projection (MIP) of the 3D volume is shown on top with depth color coded. The measured/estimated pupil wavefronts are displayed on the top right corner. The sideview MIP of the volume is shown on the bottom. Yellow arrows highlight cross sections of cells for comparison.

The plenoptic illumination encodes ocular aberrations and volume depths by recoding the propagation behavior of the beams as they are refracted and focused by the anterior ocular optics before penetrating the 3D retinal volume to excite fluorescent structures along their path (Fig1. a-ii and a-iii). In the plenoptic images, depth is represented as lateral shifts of the same structure across parallax images, which depend on both axial (depth) position and sub-pupil position (illumination angle). For demonstration, we imaged a living *Cx3cr1-GFP* mouse which labels mononuclear phagocytes, which includes microglia in healthy eyes and monocyte-derived macrophages in eyes. Figure 1c shows images of GFP-labeled retinal microglia captured under two sub-pupil positions. The cells exhibit different lateral translations between the two views, corresponding to their axial positions (Supplementary video 1). Under ideal, unaberrated optics, the illumination angle is solely determined by the sub-aperture position, and the depth of 3D structures can be recovered from the lateral shifts across plenopic views (Fig1. a-ii and b). In aberrated optics such as the eye, however, the actual incident angle of sub-pupil beams will be deviated (Fig1. a-iii and b). This aberration-induced angular deviation shifts the PSF in the retina and transversely translates the entire parallax image toward same direction regardless of depth, with magnitude and orientation determined by the local phase gradients on the pupil. Based on the depth-dependent parallax and aberration-induced shifts, they can be disentangled, the 3D retinal volume and pupil aberrations can be jointly inferred from the plenoptic images^42,43^.

### Space-varying ocular aberrations corrected with synthetic Hartmann-Shack wavefront sensor (syn-HSWS) and digital aberration correction (DAC)

In principle, the 3D volume can be computationally reconstructed from the PI-SLO images using a 3D reconstruction algorithm based on Richardson-Lucy deconvolution (Fig. 1e, Fig. S3), as demonstrated in microscope-based LFM^42,43,46^. However, even though aberration information is embedded, 3D PSF calibration using microspheres across the imaging space is required in classic LFM to robustly solve the ill-posed lightfield reconstruction problem^42–44^, which is impossible in the living eye. To calibrate the forward model PSFs of the unknown ocular optics, we implemented a synthetic Hartmann-Shack wavefront sensor (syn-HSWS) to measure ocular aberrations. The syn-HSWS deployed a PSF camera at a de-scanned detection plane conjugated to the retina to image the reflectance PSFs of the excitation beams under each sub-pupil position (Fig. 1d, Fig.S2). The sub-pupil illumination PSFs were tiled corresponding to their sub-pupil positions to create a spotgram conceptually equivalent to a conventional Hartmann-Shack wavefront sensor ^47^. As mentioned above, sub-pupil PSFs shift laterally by the local pupil phase gradient which can be measured to calculate the full pupil wavefront and construct a complex pupil function of the eye. The complex pupil function was used to provide initial calibration of 3D PSFs in the forward model for volume reconstruction as well as digital aberration correction (DAC, Fig. 1e).

The feasibility of syn-HSWS and DAC was demonstrated *in vivo* in a *Cx3cr1-GFP* mouse eye using laser-induced CNV model^48^ at two weeks following the laser injury (Fig. 1f, Fig. S8). In this model, laser damage to the retinal pigment epithelium (RPE) and Bruch’s membrane produces strong background fluorescence from the highly aggregating fluorescent cells underlying choroid, posing an additional challenge for computational reconstruction under severe ocular aberrations. Without prior pupil aberrations by syn-HSWS, the reconstruction failed to accurately reassign the fluorescent photons to the 3D cellular structures, resulting in severe artifacts regardless of whether DAC was applied, as DAC alone was not enough to correctly estimate the pupil wavefront. Using only syn-HSWS wavefront as prior without DAC, the retinal volume was reconstructed with axial positions of single cells identifiable, while image quality degraded toward the peripheral ROI as the space varying aberrations were not accounted for. Combining the syn-HSWS prior with patch-wise reconstruction with DAC to refine aberrations locally, the image quality at the periphery was improved, suggesting an aberration-corrected 3D reconstruction across the 15 × 15° FOV, exceeding the 3-5° isoplanatic patch size in the mouse eye ^19,28^. Moreover, the syn-HSWS showed improved reconstruction robustness by enabling faster convergence with reduced residual wavefront errors in the estimated pupil aberrations (Fig. S4).

### Up to third order sub-cellular processes of retinal microglia revealed in 3D under low laser power excitation using PI-SLO

Retinal microglia are resident immune cells that act as sentinels to constantly survey and the retina and promote homeostasis. Resident immune cells including resting microglia in the retina have dynamic, highly ramified processes for continuous surveillance of the neuroretinal parenchyma^8^. Monitoring and quantifying immune cells may enable early biomarkers for understanding progression of retinal diseases, including diabetic retinopathy, age-relate macular degeneration, and glaucoma ^49–51^. Due to their small size and weak fluorescent signal of the processes, these subcellular components have only been imaged *in vivo* previously by AO under a relatively high laser power >100-250µW^27^ or a ∼60s integration time^9^.

Here, we first validated large FOV aberration-corrected single cell imaging of our PI-SLO by imaging GFP-labelled retinal microglia in a living healthy transgenic *Cx3cr1-GFP* mouse using a <33µW laser power at the cornea, half of the lowest value reported by the previous work^9^. Spatial-varying ocular aberrations were estimated locally across the 15×15° FOV (Fig. 2a), revealing the field-dependent pattern between the adjacent 5° ROIs. With implementation of the syn-HSWS and multisite DAC, single microglia cell bodies as well as sub-cellular processes were clearly visualized across the 15° FOV, ∼9× larger than a typical mouse AOSLO^27,28^ (Fig.2b, Supplementary video 2). Three distinct cell plexi were identified from the sideview projection across a total thickness of 116µm (Fig. 2b-right). The distance from the superficial plexus (SP) to the intermediate plexus (IP) was 52µm, and that from the IP to the deep plexus (DP) was 64µm, matching the microglia distribution in the healthy mouse nerve fiber layer (NFL), inner plexiform layer (IPL), and outer plexiform layer (OPL), as previously observed with confocal AOSLO^26,27^. Figure 2c shows a representative microglial cell in the IPL, where sub-cellular processes were clearly visualized for up to the third branch order (Fig. 2c). Interestingly, the 3D organization of processes in the same cell was observed *in vivo* for the first time showing an axial spread of ∼36µm, suggesting the process may extend to the adjacent retinal layers such as the ganglion cell layer (GCL) or the inner nuclear layer (INL).

**Figure 2.**
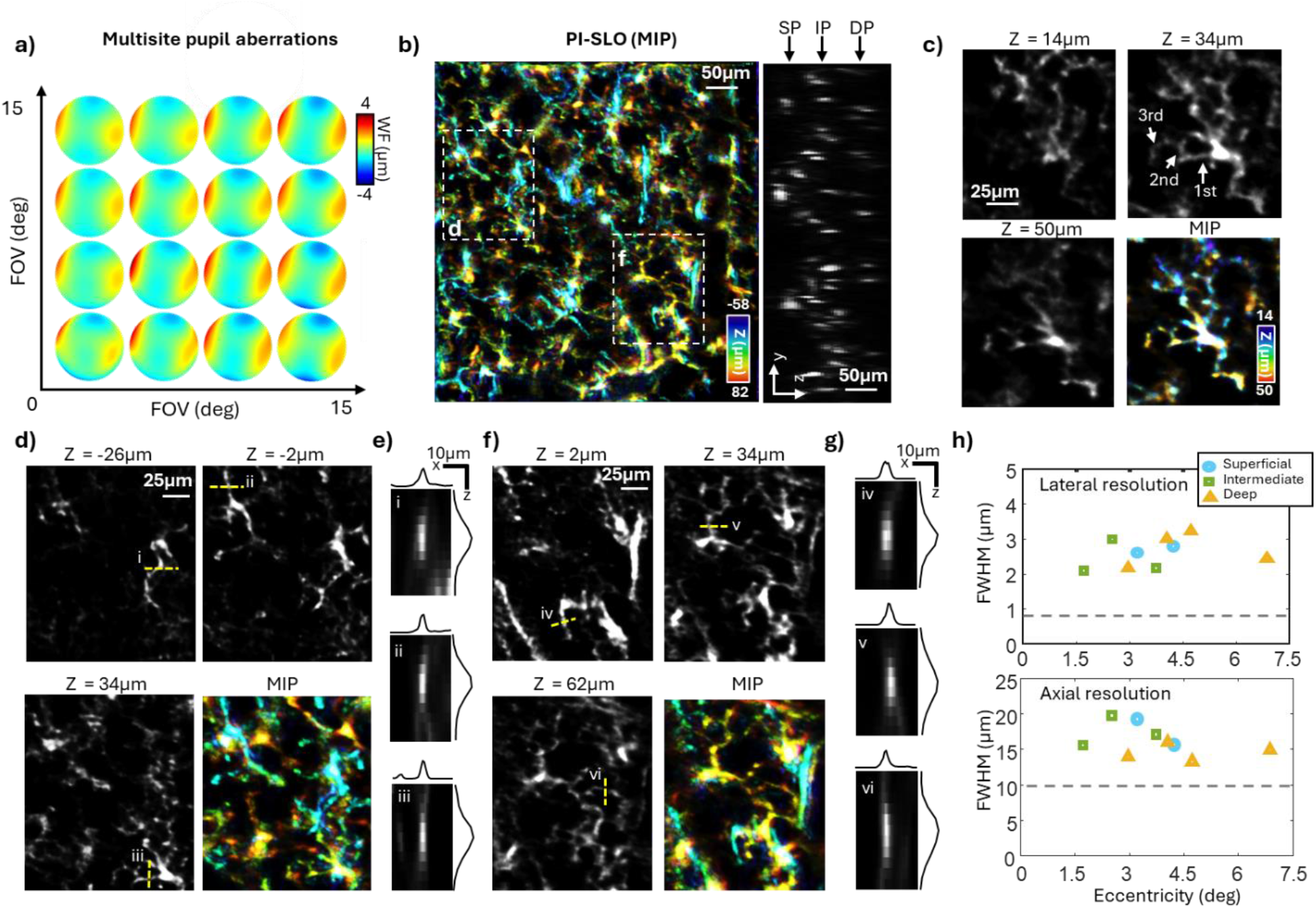
*In vivo* 3D imaging of GFP labeled microglial using PI-SLO. **a)** Estimated local wavefront map of the eye’s pupil across the 15×15 degrees FOV. **b)** Maximum intensity projection (MIP) of the reconstructed aberration-corrected 3D retinal volume. Depth is color-coded from −90 to 65 µm, where 0 µm corresponds to the native focal plane. A side-view projection is shown on the right, highlighting the axial distribution of three plexuses of microglial cells. **c)** Zoom-in image of a single microglia cell. MIP and three selected z-slices are shown. Arrows indicate visualization of up to third-order microglial processes. **d)** and **f)** Two selected zoom-in regions as labeled in **b). e, g)** Line spread functions (LSFs) of microglial processes, as indicated by the yellow dashed lines in **d)** and **f). h)** Lateral (top) and axial (bottom) resolution characterized from 9 processes selected across different lateral and axial positions, gray dashed line shows measured resolution of AOSLO previously reported^28^.

We used microglial cellular processes as the benchmark to characterize the 3D resolution of our systems in the living eye by evaluating the line spread function (LSF), as the line-shaped microglia processes have a thickness of 1-2µm^33^. Nine processes were manually selected across the 15×15° FOV. A relatively consistent LSF was observed across the full field and three cell plexuses suggesting the feasibility of 3D reconstruction with space-varying aberration corrected (Fig. 2e-g). Figure 2h shows the resolutions across the full FOV which were 2.62 ± 0.42µm lateral and 16.19± 2.20µm (Mean ± STD, N = 9 processes).

### PI-SLO simultaneously monitored motility of one-hundred resident immune cells across three retinal plexuses for over 13 minutes

Imaging the retina across an extended FOV in 3D enables simultaneous monitoring of a large scale of cells across different retinal layers. Next, we demonstrate this capability by imaging dynamics of resident immune cells across a 22° × 15° FOV again in a healthy, living *Cx3cr1-GFP* mouse eye. The high efficiency and low laser power enabled continuous imaging for over 13 minutes (800 seconds) without obvious signal or resolution degrade. The measured pupil aberrations were evaluated showing less than 2.9% change of root mean square (RMS) wavefront across 800 seconds, which suggests the stable optical quality and eye placement over the extended period of imaging time (Fig. 3b). Timelapse imaging revealed the spontaneous motility of cells with subcellular processes clearly visualized across all three layers (Fig. 3a, Supplementary video 3). A total of 93 immune cells were identified across the 22° FOV, with 22 cells in the superficial layer, 34 cells in the intermediate layer, and 37 cells in the deep layer (Fig. S5). Cell motility of different retinal layers were readily observed, including transient extensions and retractions of cellular processes (Fig. 3c), consistent with prior observations using AOSLO^9^, yet the FOV and cell numbers were 12× larger by PI-SLO. The dynamics of all 93 cells were characterized by evaluating the correlation coefficient of individual cell between the first and each point (Fig. 3d). The correlation coefficients of superficial immune cells were significantly lower than the intermediate and deep cells at the later time points. This suggested a depth dependent pattern showing cells located in the superficial retinal layer exhibited more active dynamics than those in the intermediate and deep layers. This observation was likely attributable to differences in GFP-expressing immune cell populations across retinal layers, as more motile cells such as macrophages and hyalocytes were located in the superficial retina or within the vitreous while the IP and DP were dominated by less active microglia.

**Figure 3.**
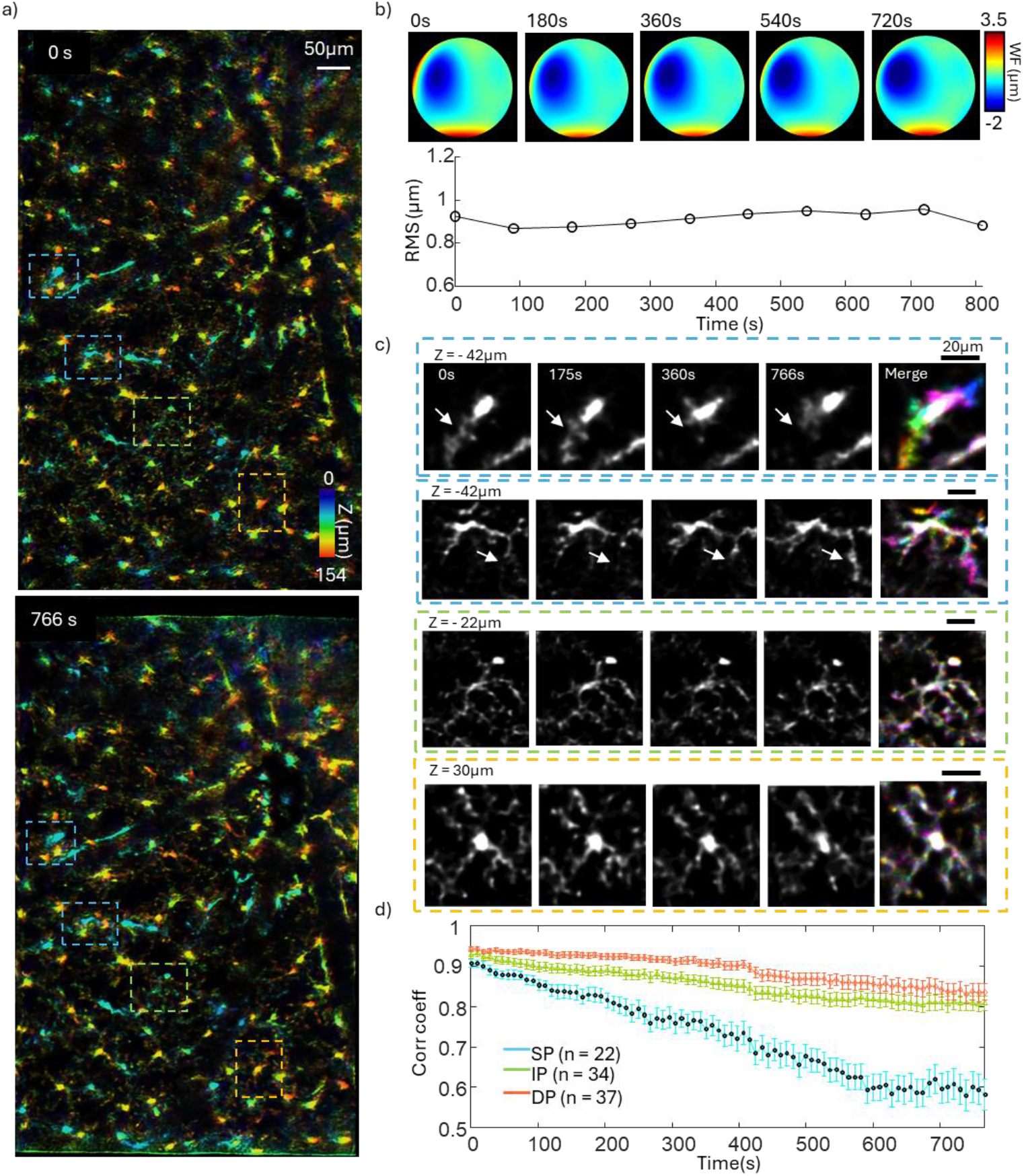
Simultaneous motility monitoring of 93 immune cells over 13 minutes. **a)** Depth color-coded MIP of reconstructed 3D images of the 0s (top) and 766s (bottom) timepoints of image acquisition. The timelapse image was captured under 15×22 degrees FOV. **b)** Estimated ocular wavefronts at representative time points. The bottom panel shows the corresponding root-mean-square (RMS) wavefront error. **d)** Quantification of motility for all 93 immune cells grouped into three retinal plexuses. Error bars: Mean ± SEM.

### Complete 3D visualization of the retinal vasculature network revealed with plenoptic illumination fluorescein angiography (FA)

Beyond single immune cells, we next applied PI-SLO to image the 3D retinal vasculature and blood flow perfusion using fluorescein angiography (FA). The inner retina contains a sophisticated 3D vascular network comprising three distinct plexi spanning from the GCL to the OPL and supporting the high metabolic demand of neural activity within resident retinal neurons. Changes in retinal blood flow perfusion and vascular morphology, including ischemia, vascular hyperpermeability, and neovascularization, are hallmarks of not only retinal diseases such as diabetic retinopathy, age-related macular degeneration, and glaucoma^52–55^, but also systemic diseases such as neurodegenerative disorders^56^.

One healthy adult mouse with retro-orbital injection of 0.1ml 2.5% FITC-dextran was imaged. The PI-SLO revealed the complete 3D retinal vasculature network from the major arteries and veins to the smallest capillaries (Fig. 4a-b, Supplementary video 4). The reconstructed volumetric images successfully resolved the retinal vascular network from large arteries and veins down to the smallest capillaries. Computational optical sectioning provided by the 3D reconstruction enabled tomographic imaging and axial localization of the three retinal vascular plexuses (Figure 4b). Notably, vertically oriented diving vessels were clearly visualized, revealing direct connections between different retinal vascular plexuses across depth (Fig. 4c). Four major types of diving vessels were identified: 1) vessels connecting all three layers; 2) vessels connecting superficial and deep layer; 3) vessels connecting intermediate and deep layer; and 4) vessels connecting superficial and intermediate layer. This observation match the morphology of mouse retinal vasculature reported previously with *ex-vivo* confocal microscopy ^57^. Visualization of these diving vessels provides a more complete representation of retinal vascular topology. In comparison, these diving vessels are often poorly resolved in clinical FA, and in optical coherence tomography angiography (OCT-A)^58^.

**Figure 4.**
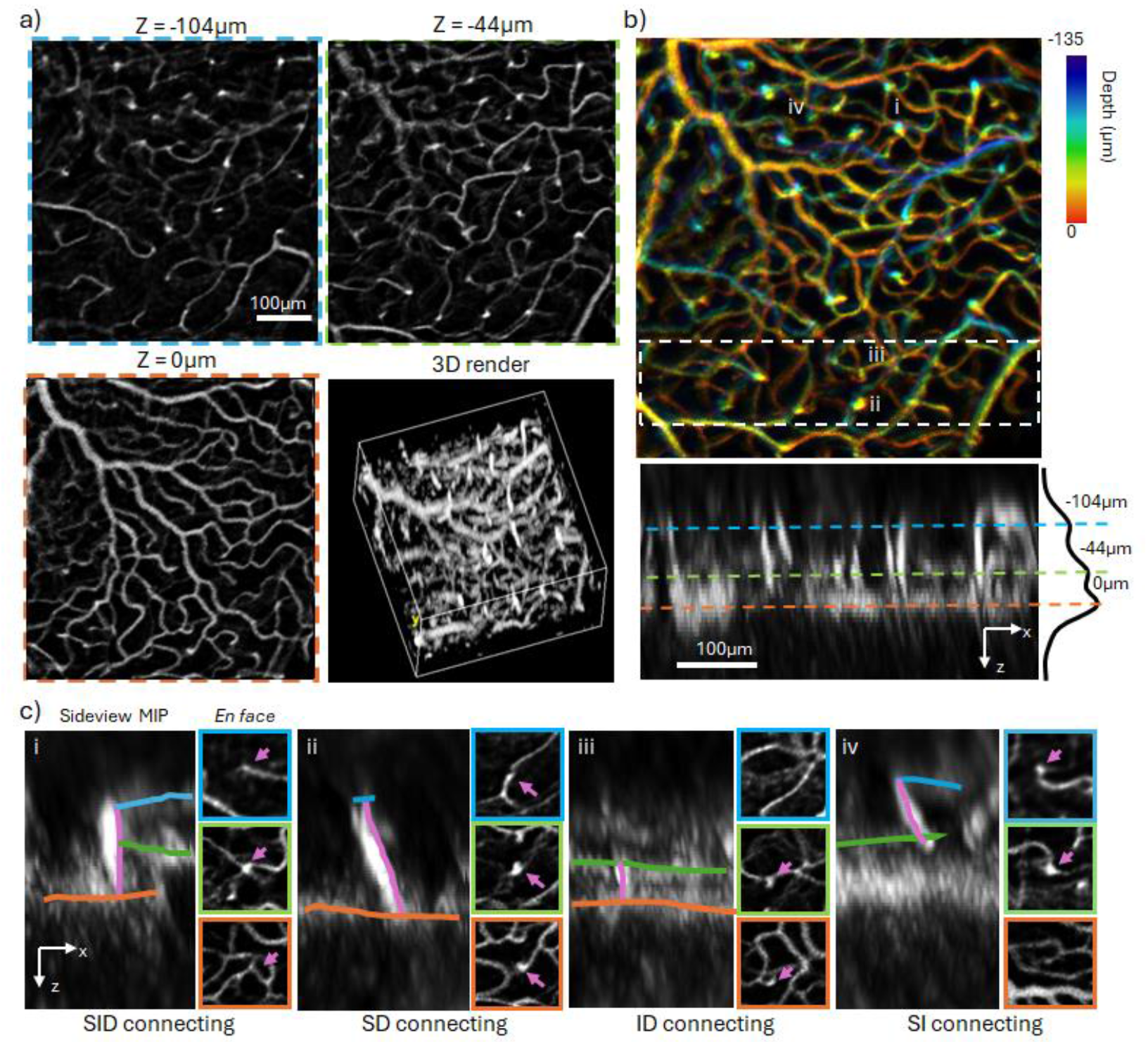
3D retinal vasculature network reconstruction with plenoptic fluorescein angiography. **a)** *En face* views of the three major retinal vascular plexuses. Bottom right: 3D rendering of the reconstructed retinal vasculature volume. **b)** Top: Depth color-coded maximum intensity projection (MIP) of the 3D vascular volume. Bottom: Side-view MIP of the region indicated by the white box in the top panel. The mean fluorescence intensity profile across depth is shown on the right, with three distinct peaks indicating the three vascular plexuses. **c)** Four representative regions illustrating distinct types of diving vessels. Three-dimensional vascular connections were manually traced and overlaid on the side-view MIPs. *En face* views of the three vascular plexuses are shown to the right of each panel, with diving vessels highlighted by magenta arrows. The locations of the four selected regions are indicated in **b)**.

### PI-SLO reveals 3D structure and calcium functions of single inner retinal neurons

Next, we investigated neuronal functions by imaging calcium dynamics in inner retinal neurons of GCaMP-expressing mice. The retina exhibits a highly stratified laminar architecture with each layer containing distinct neuron subtypes and structures, making it possible to investigate the individual neural hierarchy with volumetric imaging ^6,11,13,59^. Simultaneous imaging of neural activity across multiple retinal layers would thus enable direct interrogation of synaptic connectivity within distinct retinal circuits, advancing our understanding of fundamental visual processing and informing emerging vision-restoration strategies, such as optogenetic reactivation^60^ and retinal neuron transplantation^61^.

Wildtype C57BL6/J mice were imaged 4 weeks after intravitreal injection of an AAV vector driving CAG-mediated expression of the genetically encoded calcium indicator GCaMP6s inside inner retinal neurons, with cytosolic Ca^2+^ level reported by green fluorescence intensity^61,62^. We first imaged a living retina with 15 ×15º FOV under continuous illumination of 488nm excitation laser to demonstrate structural imaging of inner retinal neurons. Figure 5b-c shows the structural volume revealing two distinct plexuses separated by ∼50 µm in depth. Cells observed in the superficial layer exhibited a larger soma area of 156.7 ± 77.4 µm^2^ (Mean ± SD., n = 21cells), consistent with retinal ganglion cells (RGCs)^63^, whereas the mosaic structure in deeper layers were smaller and more consistent in size (71.6 ± 14.5 µm^2^, n = 27 cells), presumably axonal terminals or somata from bipolar cells (BCs)^64^. 3D subcellular structures in a single RGC, including axons and dendrites, were identified (Fig. 5c, Supplementary video 5). Axons appeared as elongated fibers in the superficial layer extending toward the optic disc, while dendrites formed radial arbors surrounding the somata extended toward deeper layers. Notably, PI-SLO reconstruction also revealed stratification of RGC dendrites (Fig. S6). This laminar dendritic structure is an important morphological parameter associated with RGC stimulus feature selectivity and subtype ^65^, which has been previously visualized *in vivo* only with AOSLO^28^.

**Figure 5.**
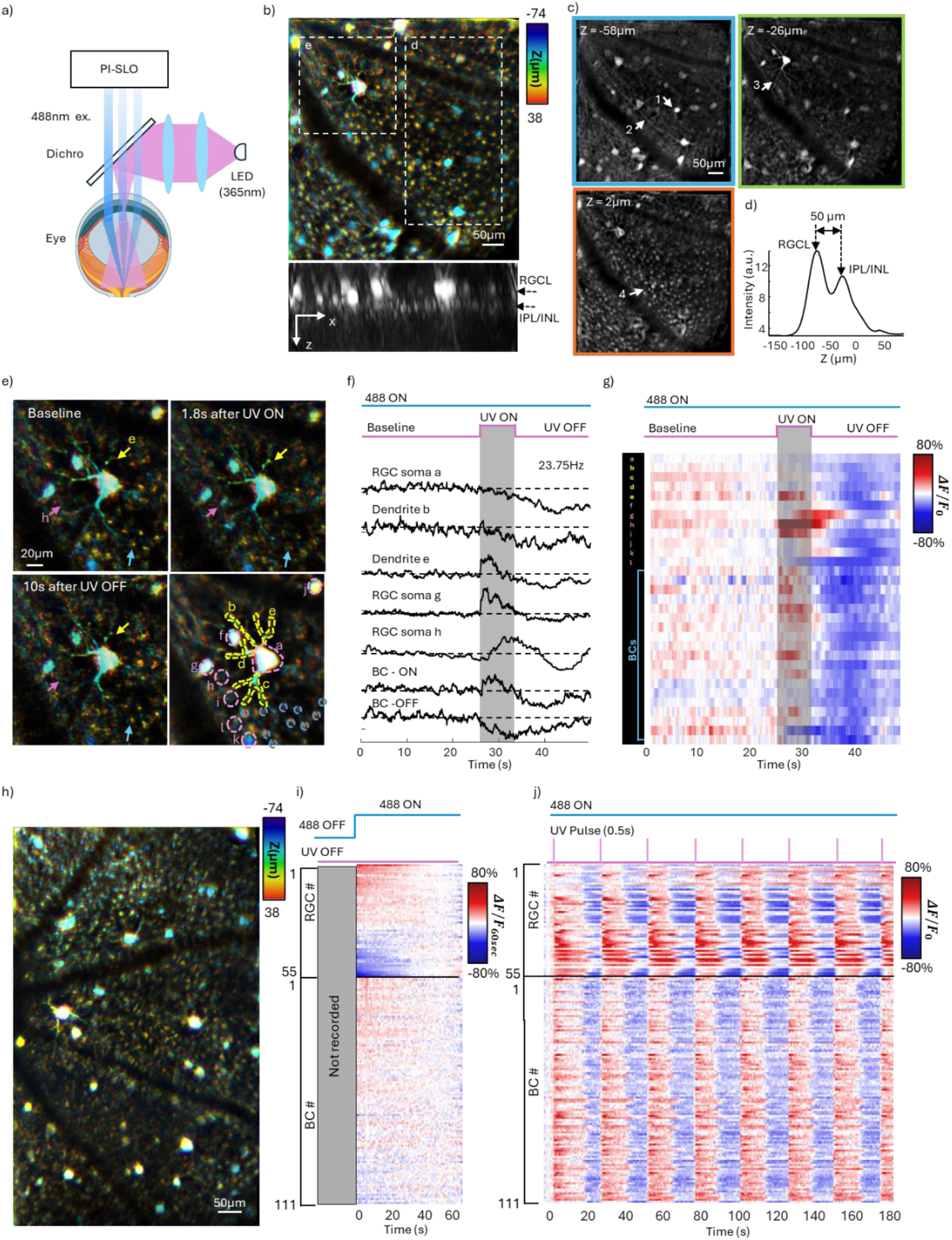
Volumetric calcium imaging in a living mouse eye. **a)** Schematic of the experimental setup. An ultraviolet (UV, 365nm) LED was deployed to provide full-field visual stimulation across the whole retina. **b)** Top: Depth color-coded maximum intensity projection (MIP) of the 3D structural image of retinal neurons. Bottom: Side-view MIP revealing two distinct neuronal layers. **c)** Three representative axial slices from the structural volume. Arrows highlight distinct cellular and subcellular features: (1) retinal ganglion cell (RGC) soma, (2) RGC axon, (3) RGC dendrites, and (4) bipolar cell (BC). **d)** Mean fluorescence intensity as a function of depth, evaluated from the boxed region in (b). Two distinct peaks indicate an axial separation of 50 µm between two neuronal plexuses. **e)** Time-lapse depth color-coded MIP of a zoomed-in region centered on a large RGC. Arrows highlight three structures exhibiting calcium responses. Bottom right: 21 manually defined regions of interest (ROIs) used for signal extraction. Label color indicates structure type: magenta, RGC soma; yellow, RGC dendrite; cyan, BC. **f)** Extracted calcium traces from six representative structures, showing distinct responses to UV stimulation. **g)** Heatmap of calcium activity from all 21 ROIs in response to UV stimulation. **h)** Structural image covering a 15 × 22-degree field of view used for large-scale analysis. **i)** Calcium responses from 222 neurons (99 RGCs and 123 BCs) immediately after onset of the 488-nm imaging laser. Most cells showed saturation in approximately 60 s after exposure. **j)** Calcium responses from the same 222 neurons in response to repeated 0.5-s UV stimulation pulses.

Calcium responses to visual stimulation were evaluated using 365-nm ultraviolet (UV) light, which selectively stimulates S-opsins in mouse cone photoreceptors^6^. Imaging was recorded for 47s with an 8-s UV stimulation following a 26-second baseline recording (Fig. 5e, Supplementary video 6). To maximize temporal resolution, we performed a rolling window reconstruction to achieve a volume rate up to 23.75 Hz, as each angular frame contains transient temporal information of the entire 3D volume^43^. Figure 5f–g shows calcium signals extracted from different neuronal structures. Distinct stimulus-evoked responses were observed across cells, with both ON and OFF responses detected in RGCs and BCs, consistent with subtype-specific functional organization^6,12,13^. Notably, distinct calcium dynamics exhibited in dendrites in the same RGC, differing not only in temporal profiles but also in response polarity, suggesting compartmentalized signal integration within single neurons^25,66^.

### PI-SLO enables monitoring of calcium signal firing from two retinal layers simultaneously across over 22°

Finally, we performed a large-scale imaging by expanding the FOV to 15×22º (Fig. 5h). Neuronal responses to 488 nm excitation laser were first evaluated. A 60s timelapse image recording was initiated a few seconds before laser onset after ∼5 min of dark adaptation. In total, 166 cells were manually identified with 55 RGCs and 111 bipolar cell soma/terminals by an experienced examiner (Fig. S7). Figure 5i shows calcium signals extracted from all 166 cells, exhibiting either hyperpolarizing or depolarizing responses upon illumination onset in both RGCs and BCs suggesting ON and OFF responses towards the excitation light (Supplementary video 7). Fluorescent traces were evaluated with normalization to the last time point when signal saturated (t=70s). All selected cells reached equilibrium within 50s without noticing obvious high frequency fluctuation, consistent with previous reports^6,12^. This suggests that sub-pupil scanning plenoptic illumination and the raster scanning SLO would not introduce confounds of measurable functional perturbations upon later UV stimuli. We next recorded calcium responses within the same FOV using 7 trials of 0.5-second ultraviolet (UV) stimulation pulses in 180 seconds (Supplementary video 8). Figure 5j shows the extracted fluorescent calcium traces from all cells. Individual neurons exhibited reproducible calcium transients in response to each stimulus pulse. Traces in Fig. 5j are presented in the same order as in Fig. 5g for direct comparison, revealing most neurons respond to UV stimulation despite being saturated from the 488nm excitation beam. To the best of our knowledge, this is the first time to achieve volumetric calcium dynamic imaging *in vivo* in the retina with two retinal layers simultaneously recorded.

## DISCUSSION

We present a computational ophthalmoscopy approach, PI-SLO, that enables single-cell–resolved 3D fluorescent imaging in the living mouse eye. Plenoptic illumination imaging captures the retinal volume under multiple angular parallax views, enabling simultaneous monitoring of cellular morphology, motility, and functional activity across multiple retinal layers without sequential refocusing, which suffers from slow acquisition speed and temporal offsets between axial planes. With syn-HSWS to measure the severe pupil wavefront, ocular aberrations were digitally corrected across the large FOV with DAC without using complex and expensive hardware adaptive optics. Together, PI-SLO establishes a practical platform for high-throughput, longitudinal interrogation of retinal single cell dynamics *in vivo*, enabling studies of retinal physiology and pathology across layers that were previously challenging.

Like other computational imaging strategies, PI-SLO requires an accurate forward model that accounts for the strong and spatially varying ocular aberrations. The syn-HSWS played an essential role by adaptively pre-calibrating forward model the strong and uncertain aberrated ocular optics *in situ* for robust 3D reconstruction. In contrast, although plenoptic measurements in some LFM technology encode aberrations that can be computationally corrected, pre-calibration by imaging standard targets (for example, fluorescent beads) is typically necessary to characterize the system PSFs ^40,42,43^, which is impossible for *in vivo* retinal imaging. Further implementation of patch-wise DAC locally fine-tuned space-varying aberrations, permitting a single cell level 3D resolution across a FOV up to 15×22º, >12× larger than a typical mouse AOSLO^25,27,28^, beyond isoplanatic limit of the mouse eye. This substantially increases imaging throughput by allowing simultaneous observation of large cell populations without image montaging. In theory, the only limiting factors of the maximum FOV are the optical design and the scanning range of the SLO system. A future development can optimize the SLO optical design and deploy a large angle optical scanner to expand the FOV comparable to state-of-the-art wide-field SLO systems, over 50–100° (1.7-3.4 mm in mouse eyes), opening the possibility of mesoscale 3D single-cell fluorescence imaging in the living eye non-invasively.

Correcting ocular aberrations allows for precisely ranging fluorescent structures in the retinal volume by exploiting the maximum angular disparity of the full pupil, enabling high resolution 3D reconstruction without AO. We demonstrated a measured axial resolution of ∼16 µm, close to the ∼11 µm measured in AOSLO ^28^ and >4× better than a non-AO cSLO^16^. The lateral resolution and depth-of-field (DOF) in PI-SLO can be flexibly adjusted by changing the sub-pupil size on the DMD to balance high resolution with extended depth of field to match experimental needs. In this work, we demonstrated a lateral resolution of 2.6 µm measured in the *in vivo* mouse retina while achieving a >100 µm DOF. This lateral and axial resolution in PI-SLO was sufficient to visualize many fine 3D cellular and sub-cellular morphology over a large FOV, including microglia processes, small capillaries, RGC dendrites, and the bipolar cell mosaic, although being ∼3× worse than the near diffraction-limited AOSLO. These details could only be resolved *in vivo* previously using AOSLO, wherein the small FOV limited cell throughput to only one or a few cells imaged in a time. While the reported absolute depth was based on assumption of previously reported ocular optical properties (2.6 mm focal length and refractive index of 1.33)^19^, the reported depths show agreement with the known anatomical organization of the mouse retina. Moreover, AOSLO typically requires a relatively high excitation power (>200µW) or a prolong integration time (>10s) to resolve such fine sub-cellular structures given the very weak fluorescent signal ^9,27,28,37^. Because the emitted fluorescent photons are captured from the entire volume without the exclusion of a confocal pinhole, the photon collection efficiency is maximized in PI-SLO. The optimized photon collection efficiency permits imaging of the same single cells with a laser power 2-8 times lower than an AOSLO and without requiring frame integration^9,27,28^. This low-power, high-efficiency characteristic is particularly advantageous for *in vivo* retinal imaging by reducing phototoxicity and enabling prolonged continuous observation of cellular dynamics.

Together, PI-SLO has enabled simultaneous imaging of the single retinal cells across a large FOV under low excitation power. Because the entire retinal volume is captured simultaneously, PI-SLO enables ‘true’ 4D (volume + time) imaging of cellular dynamics and functional readout without time offset confound between different axial planes. In the current setup, the plenoptic illumination imaging captures the angular parallax images under 23 frames per second. Because the entire 3D volume is projected in each individual parallax image, each angular image contains the transient temporal information of the volume, enabling a 23 Hz volume rate with rolling window reconstruction. Similar rolling window reconstruction has been widely implemented to enhance temporal resolution in other modalities using multiple acquisition, such as computed tomography and computational microscopy ^42,44^. The fast transient cellular dynamics, such as light-evoked calcium fluxes within neurons, can now be captured in 3D. Future work can further improve the volumetric rate for *in vivo* functional imaging with fast sensors, such as those detecting transmembrane voltage and glutamate transmission ^67–69^. The volume rate can be improved by simply using a faster optical scanner, such as microelectromechanical systems (MEMS) scanners^70^ or polygon mirror^71^. Additionally, reducing the FOV or implementing optimized scanning protocols, such as sparse sampling^72^ or random-access scanning^73^, could further enhance temporal resolution when required.

To the best of our knowledge, we attained the first 4D calcium imaging of the living mouse retina by simultaneously capturing calcium signals from RGCs and BCs within two distinct retinal layers. This capability is particularly valuable for studying interactions across hierarchical retinal layers, particularly to characterize neural signal transduction along the visual pathway. Thanks to high-resolution provided by syn-HSWS /DAC and maximized detection efficiency provided by non-confocal detection, RGCs somata as well as their dendritic arbors —despite weak fluorescence— were easily identified with GCaMP. The mosaic structures in the deeper layer were attributed to BCs (either axon terminals or soma) depending on their size, arrangement, and axial locations, while future works are required to validate with *ex vivo* histology to identify which compartments in these interneurons were labelled. The depth resolving capability of our approach provides opportunity to simultaneously image the signal transduction through different neuron types using the same fluorescent reporter/excitation line. A future direction will ubiquitously labeled different retinal neuron types, including photoreceptors, enabling optical readouts of neural connectivity and signal transduction across full retinal circuits. More complex stimulation protocols—including single-photoreceptor targeting, spatially patterned stimuli, and temporal chirp sequences—can further dissect subtype-specific responses and functional connectivity *in vivo*. Notably, the point-scanning architecture of our PI-SLO is naturally compatible with two-photon excitation using near-infrared illumination ^43^, allowing simultaneous visible-light visual stimulation.

Overall, PI-SLO provides a large-field, single-cell–resolved *in vivo* 4D fluorescent imaging platform that integrates structural and functional imaging without adaptive optics hardware, paving the way toward transforming the living eye into a versatile *in vivo* window for studying retinal and central nervous system physiology and disease.

## ONLINE METHODS

### System design of plenoptic illumination scanning light ophthalmoscope (PI-SLO)

The schematic of our PI-SLO system is shown in Figure. S1. Our LF-SLO system built upon a lens-based SLO designed for mouse eye with a 2mm entrance pupil. The 488nm excitation beam emitted from a laser diode (SAPPHIRE 488-50, Coherent, PA, USA) was collimated and expanded to a 10 mm diameter beam size. The collimated beam was delivered to the digital micromirror device (DMD, DLP® LightCrafter™ 4500, Texas Instruments, TX, USA), where micromirror facets in the ON state reflected the zeroth-order, undiffracted light into the downstream optical path. Light from OFF state facets, as well as higher-order diffracted components, was rejected by an aperture stop to ensure clean illumination. The DMD was placed conjugate to the pupil of the eye. The effective pupil area corresponded to a 4 mm diameter region on the DMD, occupying less than half of the 10 mm collimated beam profile. This configuration ensured less than 50% intensity variation across different sub-pupil regions. The beam was then delivered through a cascade of five 4-f afocal telescopes into the eye, where the DMD was at the 1^st^ pupil plane and the eye was at the 6^th^ pupil plane. To provide raster scanning, the beam was steered by two scanning mirrors deployed at the 3^rd^ and 4^th^ conjugate pupil planes, respectively. The fast axis was controlled by a resonant scanner (SC30, EOPC, CA, USA) with 10kHz line rate, while the slow axis was a galvanometer (GVS011, Thorlabs, NJ, USA). The maximal scanning optical angle in the eye was 15º on the fast axis and 22º on the slow axis. An electric tunable lens (ETL, EL-3-10-VIS-26D-FPC, Optotune AG, Dietikon, Switzerland) was deployed at the 5^th^ pupil plane close to the eye to adjust the native focal plane on the retina. The emitted fluorescent photons returned from the retina were collected through the full pupil of the eye. The fluorescent light delivered through the common path before being reflected into the detection path by a dichroic mirror (RT488rdc, Chroma Technology Corp., VT, USA) after L13. Another 4-f afocal telescope was implemented to demagnified and imaged the fluorescent light from the pupil into a photomultiplier tube (PMT, PMT210, Thorlabs, NJ, USA). Another iris aperture stop was placed at the conjugate image plane. The aperture diameter was carefully adjusted to collect fluorescent signal from the retinal volume, while suppressing autofluorescence from the contact lens and retinal pigmented epithelium induced by the UV stimulation. The fluorescence signal detected by the photomultiplier tube (PMT) was digitized using a high-speed digitizer (ATS9440, Alazar Technologies Inc., QC, Canada) for computer acquisition.

### Image acquisition and preprocessing

To realize plenoptic illumination imaging, the scanners, DMD, and the digitizer were synchronized with a data acquisition card (DAQ, DAQ 6341, National Instruments, TX, USA). The TTL signal output from the resonant scanner was used to synchronize the digitizer for line acquisition (H-sync). Meanwhile, the DAQ card was also synchronized by the H-sync to output two waveforms to control the galvonmeter for slow axis scanning, and the digitizer for frame acquisition (V-sync). The DMD was synchronized by the V-sync signal to switch pattern display to ensure each frame was acquired under a specific illumination sub-pupil. The binary DMD patterns were preloaded into the controller. In this work, a total of 20 sub-pupils (patterns) were used (Fig. 1S). The sub-pupil diameter was carefully selected to be 320µm in the eye to balance the depth of field and lateral resolution for imaging single cells in the inner retina. The SLO captured 2D images under a 23.75 frames per second frame rate, while the time to complete a plenoptic acquisition was 0.85 seconds.

Because the resonant scanner scanned the beam via a sinusoidal pattern, preprocessing has to be done to rectify the image into a square pixel grid. The image was first desinusoided with a cross-correlation based algorithm previously presented by Yang et al ^74^. The pixel pitches on the fast and slow axes were pre-calibrated by imaging a Ronchi ruling with a stand achromat lens, allowing 2D interpolation to rectify the image into square pixel. One-time Ronchi ruling calibration is needed only after the system is realigned. The plenoptic image sequence was generated by indexing each image with corresponding DMD pattern for computational reconstruction.

### Aberration measurement using synthetic Hartmann-Shack wavefront sensor (syn-HSWS)

The synthetic Hartmann-Shack wavefront sensing was implemented by imaging the de-scanned PSF reflected from the retina (Fig. S1). To do this, a polarizing beamsplitter (PBS 201, Thorlabs, NJ, USA) was implemented between the dichroic mirror and L2 to allow the excitation light to transmit through while reflecting the returned scattered light from the retina. The returned scattered light was focused by lens (L14) and imaged by a CMOS camera (DCC1240M, Thorlabs, NJ, USA). A linear polarizer was deployed in front of L14 to suppress specular reflection from the lens elements in the system. A quarter waveplate (WPQ20ME-488, Thorlabs, NJ, USA) was finely tuned to maximize intensity of imaged PSF. The camera frame acquisition was also synchronized to V-sync signal, allowing it to capture individual PSF profile under specific sub-pupil illumination. The exposure time and gain were carefully adjusted to integrate as much as the scanning time for a SLO frame acquisition, while not saturating camera pixels.

The PSF images were tiled according to the DMD sub-pupil pattern to generate the syn-HSWS spotgram (Fig. S2). Centroiding of each PSF was measured by calculating the pixel coordinate of the max intensity. Shifts induced by local phase gradients were evaluated by comparing each PSF centroid with reference origin of a flat wavefront (Fig. S2b). To calibrate a flat wavefront reference origin, a paper was placed at the focus between L3 and L4. The flat wavefront reference was determined by measuring the centroids of PSFs reflected from the paper. Similar to a conventional MLA-based HSWS, the Zernike coefficients of the eye’s pupil wavefront *c* were calculated from the PSF shifts ***S***(*i*) = (*S*_*x*_(*i*), *S*_*y*_(*i*)) by straightforward Zernike fitting ^47^:

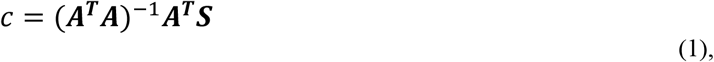

where *i* denotes the index of sub-pupil, ***A***(*i, p*) = (*A*_*x*_(*i, p*), *A*_*y*_(*j, p*)) describes the Zernike gradient matrix:

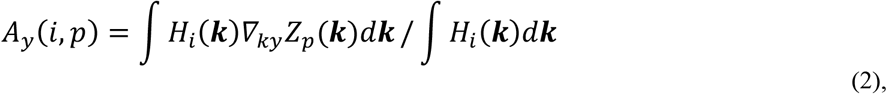

And

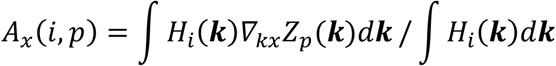

where ***k*** = (*kx, ky*) represents the coordinate of pupil plane, *Z*_*p*_(***k***) denotes *p*th order 2D Zernike polynomial, while *H*_*i*_ (***k***) denotes the *i*th sub-pupil aperture pattern. The pupil wavefront *WF*(***k***) was finally reconstructed as (Fig. S2c):

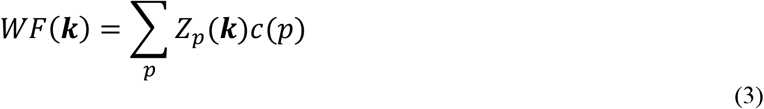

In this work, we reconstructed the wavefront with 17 Zernike orders which is enough to describe the wavefront of mouse eye, which is dominated by the lower-order aberrations up to 13th Zernike orders ^19^. The syn-HSWS was validated by imaging with a standard achromatic doublet, a 3mm diameter ball lens, and the *in vivo* mouse eye (Fig. S2).

### Jointly estimation of space varying ocular aberrations and 3D retinal volume

The 3D fluorescent volume was computationally reconstructed from the acquired space-angle images by a custom software written in Matllab. The algorithm jointly reconstruct the 3D retinal volume and estimate the ocular aberrations based on a Richardson-Lucy deconvolution inspired by the previous work in lightfield microscopy ^42,46,75^. Figure S4 shows the reconstruction routine. The reconstruction was performed patch-wise to address the space-varying aberrations in the mouse eye. The full FOV plenoptic images were first partitioned into 5º patches, matching the isoplanatic patch size of mouse eyes as described in previous works with AOSLO^11,76^. To provide seamless merging after reconstruction, XX% overlaps were applied when segmenting the patches.

The 3D reconstruction and digital aberration correction (DAC) were performed patch-by-patch. To initialize the reconstruction, the pupil aberrations measured by the syn-HSWS *WF*(***k***)were used to estimate the initial 3D PSFs for the forward model. The complex pupil function of the eye was first constructed: *P*(***k***) = |*P*(***k***)|*e*^−*iWF*(***k***)^, where the magnitude |*P*(***k***)| was determined by the transmission distribution of the eye and the laser power of each sub pupil illumination, which can be calculated by up sampling the corresponding spatial distribution of mean image intensities on the pupil. The estimated 3D PSFs *h*_*i*_(*r, z*) were calculated from pupil function *P*(***k***) and individual DMD sub-pupil patterns *H*_*i*_ (***k***) using angular spectrum propagation^77^:

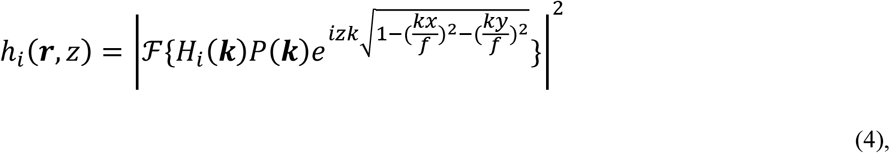

Where ℱ{. } represent 2D Fourier transform, ***r*** = (*x, y*) denotes the transvers coordinate of the retinal space, *z* denotes depth in the retina volume, *f* denotes the focal length of the mouse eye which assumed to be 2400µm for inner retinal imaging in this work^19^, *k* = 2*πn*/λ denotes wave vector was calculated from the wavelength λ = 0.488μm and refractive index of the retina/vitreous *n* which is assumed to be 1.33. The forward model of plenoptic illumination imaging *I*_*i*_ (***r***) was described as:

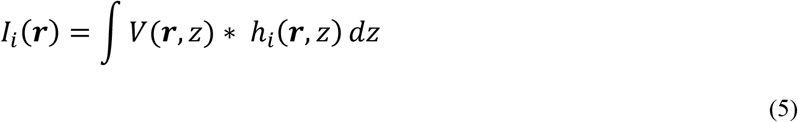

Where {*} denotes 2D convolution, *V*(***r***, *z*) denotes the 3D fluorescent retinal volume.

A Richardson-Lucy deconvolution was deployed to update the 3D volume by iteratively optimizing the Poisson likelihood 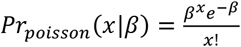 between the measurement and the estimation ^46,78,79^:

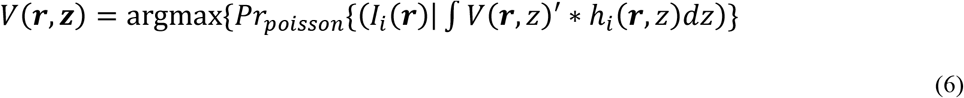

The estimated volume was updated by deconvolving the measured space-angle image with the estimated 3D PSFs angle-by-angle. In this work, the z-spacing in the reconstruction volume was set to 4-6µm, less than a half of the theoretical axial resolution of the reconstruciton. After all angles visited, the estimated volume was forward propagated used to generate a sequence of simulated space-angle images. These simulated space-angle images were used as reference templates to estimate the aberration-induced image shifts by calculating the 2D cross correlation with the measured images. These shifts were linear to the corresponding phase gradient of the residual pupil wavefront errors. The residual wavefront error was calculated in the same way as syn-HSWS by using. Eq 1-3. The residual pupil wavefront errors updated the estimated pupil wavefront as well as the 3D PSFs in the next iteration to compensate the local aberrations. This process performed iteratively until convergence. Finally, the reconstructed volumes from patches were merged with a gaussian-weighted averaging in the end to construct the full field 3D image. All data presented in this work was reconstructed with 10 iterations, except for the timelapse calcium imaging. The reconstruction software was written with Matlab script running on a personal computer (CPU: Intel Core i9-11900K, RAM: 128GB DDR5, and GPU: NVIDIA GeForce 4070). The total computation time to reconstruct a volume was ∼ 10minutes.

### Animal preparation

All animal experiments were conducted in accordance with the ethical guidelines for animal care and received approval from the Institutional Animal Care and Use Committee at Johns Hopkins University. *In vivo* imaging was performed with healthy mice. All mice were anesthetized first with intraperitoneal ketamine/xylazine injection and maintain with inhalation of a mixture of 1% (v/v) isoflurane and oxygen. The pupil of the mouse eye was fully dilated with drops of 1% tropicamide and 2.5% phenylephrine. A +10D contact lens was then applied on the imaging eye for a smoother wavefront, maintaining hydration of the cornea and prevents cataract. Meanwhilem, artificial tear gel (Genteal, Alcon Laboratories Inc, Fort Worth, TX) was applied every 30 minutes during imaging to further maintain hydration for a longer imaging time. Body temperature was maintained at 37 °C using a heating pad throughout imaging.

### Characterization of lateral/axial resolution in resident immune cell images

Images presented in Fig. 2 were acquired from a healthy CX3CR1-GFP mouse, in which resident immune cells (macrophages and microglia) were labeled with green fluorescent protein. The maximum laser power used in this experiment was 33µW at the cornea. Because the dynamics of resident immune cells are relatively slow^10^,^24^, plenoptic images captured under the same illumination angle were averaged over ten consecutive sequences before reconstruction to improve the SNR and enhance visualization of fine subcellular processes. Before averaging, a rigid-registration based on 2D cross-correlation was performed to compensate eye motions for the ten sequences.

To quantify the lateral and axial resolution of the PI-SLO reconstruction, nine thin subcellular processes were manually selected from the reconstructed 3D volume for line spread function (LSF) analysis. The processes were carefully chosen by an experienced user to ensure they are laterally and axially isolated from neighboring structures. LSFs were obtained from cross-sectional intensity profiles perpendicular to the selected processes, with ten profiles averaged along each process to reduce noise. The full width at half maximum (FWHM) measured in the lateral and axial directions was used to characterize the resolution.

### Tracking resident immune cell motilities

A healthy CX3CR1-GFP mouse was used to monitor resident immune cell motility in vivo with a maximum excitation power of 33 µW at the cornea. The same region of interest (ROI) was continuously imaged for 800 s immediately following application of artificial tear gel to maintain ocular hydration throughout the imaging session. As described above, plenoptic images acquired under identical illumination conditions were registered and averaged for each sequence prior to reconstruction, yielding an effective temporal resolution of ∼0.1 Hz for monitoring immune cell dynamics. To compensate for eye motion during the 800 s recording period, the reconstructed time-lapse 3D volumes were further aligned using a demons-based nonrigid registration algorithm implemented in MATLAB. Registration was performed on a volume-by-volume basis, using the 3D volume acquired at the 400 s time point as the reference template.

To characterize motilities of individual resident immune cells, 91 rectangular ROIs containing single cells were manually selected using ROI Manager in ImageJ(https://imagej.nih.gov/ij/) from the reconstructed 3D volume at the first time point. For each ROI, the best focal plane of the cell was manually identified. Maximum intensity projections (MIPs) were then computed over an axial range of ±18 µm around the focal plane to account for the three-dimensional morphology of the cells and to mitigate small axial displacements (Fig. S5). Morphological dynamics of individual cells were quantified by calculating the cross-correlation coefficient between the ROI at the first time point and the corresponding ROIs at subsequent time points (Fig. 3d). Lower correlation coefficients indicate greater morphological activity, whereas higher coefficients correspond to more stable cellular morphology.

### Longitudinal imaging of a laser-induced choroidal neovascularization (CNV) mouse model

Laser-induced choroidal neovascularization (CNV) was performed in a healthy CX3CR1-GFP mouse. The animal was anesthetized by intraperitoneal injection of ketamine (100mg/kg) and xylazine (10mg/kg), and the pupils were dilated with a topical drop of 1% tropicamide. Lubricating eye gel (GenTeal Tears, Alcon) was applied to the cornea before fundus imaging. The fundus was visualized using an imaging camera to locate the optic disc (Fig. S6a), and laser photocoagulation was performed using an image-guided laser system (Micron III®, Phoenix-Micron, Inc., OR, USA) with the following parameters: 532 nm wavelength, laser power 240 mW, 70 mS duration, and a 50 um spot size, as previously described^80^. Three laser burn were created around the optic nerve while avoiding major retinal blood vessels. Rupture of Bruch’s membrane was confirmed by the appearance of a vaporization bubble following laser photocoagulation, indicating a successful laser burn.

Longitudinal PI-SLO imaging was performed in the eye with laser ablation. We first acquired baseline images 48 h prior to laser ablation. Longitudinal imaging sessions were subsequently conducted at 30 min post-ablation (D0), 5 days (D5), and 14 days (D14) after laser treatment (Fig. S6b). At each time point, 9–15 images with a 15° × 15° field of view (FOV) were acquired and stitched to generate a montage covering all three laser lesions (Fig. S4c). The maximum laser power for excitation 33µW at the cornea.

### Fluorescein angiography imaging

For fluorescein angiography (FA), mice were pharmacologically dilated and anesthetized with ketamine/xylazine. A 0.1 mL bolus of 2.5% fluorescein isothiocyanate–dextran (46944-500MG-F, Sigma-Aldrich) was administered via retro-orbital injection to visualize the retinal vasculature. The maximum laser power for excitation was 5µW at the cornea.

### *In vivo* calcium imaging of inner retinal neurons

*In vivo* calcium imaging was performed with healthy mice for at least 4 weeks after intravitreal injection of pAAV-CAG2-GCaMP6s. To stably induce expression of GCaMP6s in the retina of C57BL/6J mice, 1.5μL of 2.5×10^13^vg/mL pAAV-CAG2-GCaMP6s was injected into the vitreous chamber of the eye using a 10μL Hamilton syringe with 34G metal needle. Intravitreal injections were performed under anesthesia (2.5% inhalation isoflurane) and with transretinal injections 3mm posterior to the supertemporal limbus. The needle was maintained in the eye for 1min to allow for equilibration of the IOP, reduced reflux, and AAV settlement in the vitreous chamber.

An experiment was first designed to evaluate the calcium response of retinal neurons to the 488 nm excitation laser without ultraviolet stimulation. was designed to evaluate the calcium response of retinal neurons to the 488 nm excitation beam in the absence of ultraviolet (UV) visual stimulation. Prior to image acquisition, the mouse was dark-adapted for at least 3 min. Imaging began with the laser shutter closed, and the shutter was opened 3-5 s after recording started. The PI-SLO system then continuously imaged the retina for 60 s following laser onset.

To provide ultraviolet (UV) visual stimulation, a UV LED (M365L3, Thorlabs, NJ, USA) was integrated into the system to deliver visual stimuli (Fig. S1). Light emitted from the LED was focused onto the pupil by the last lens in the illumination path, producing wide-field illumination across the retina. The UV power was adjusted to ∼10 µW at the cornea. The stimulus timing was synchronized with PI-SLO frame acquisition using an additional DAQ device (USB-6003, National Instruments, TX, USA). Before imaging, the mouse retina was exposed to the scanning 488 nm excitation beam for at least 1 min before image acquisition to saturate baseline neuronal calcium activity.

### Image processing and analysis of calcium imaging

Reconstruction of time-lapse 3D volumes for in vivo calcium imaging was performed using a rolling-window approach to exploit the temporal information embedded in the 23.75 Hz 2D frame rate. To reduce computational cost and improve reconstruction signal-to-noise ratio (SNR), a high-quality structural volume was first generated by averaging a sequence of 50 plenoptic (space–angle) images. This structural volume, together with the estimated pupil aberrations, was used as the initial condition for reconstructing each time-point volume. Each volume converged within two iterations, with digital aberration correction (DAC) applied only at the initial stage to compensate for minor phase variations and eye motion. The reconstruction time for a single time-point volume was approximately 40 s. Timestamp of each frame was provided by the PI-SLO system.

For calcium signal extraction, neuronal somata (and synaptic regions shown in Fig. 5e–g) were manually identified and segmented by an experienced examiner using ROI Manager in ImageJ(https://imagej.nih.gov/ij/). A script written in Matlab was used to calculate the fluorescence intensity by averaging pixel values within each segmented region, and cell types were classified based on morphology, size, and depth. Background fluctuations arising from autofluorescence of the contact lens and retinal pigment epithelium (RPE) during UV stimulation were estimated using signals from manually segmented large blood vessel regions^12^.

