## Supplementary Information for "Computational aberration-corrected volumetric imaging of single retinal cells in the living eye"

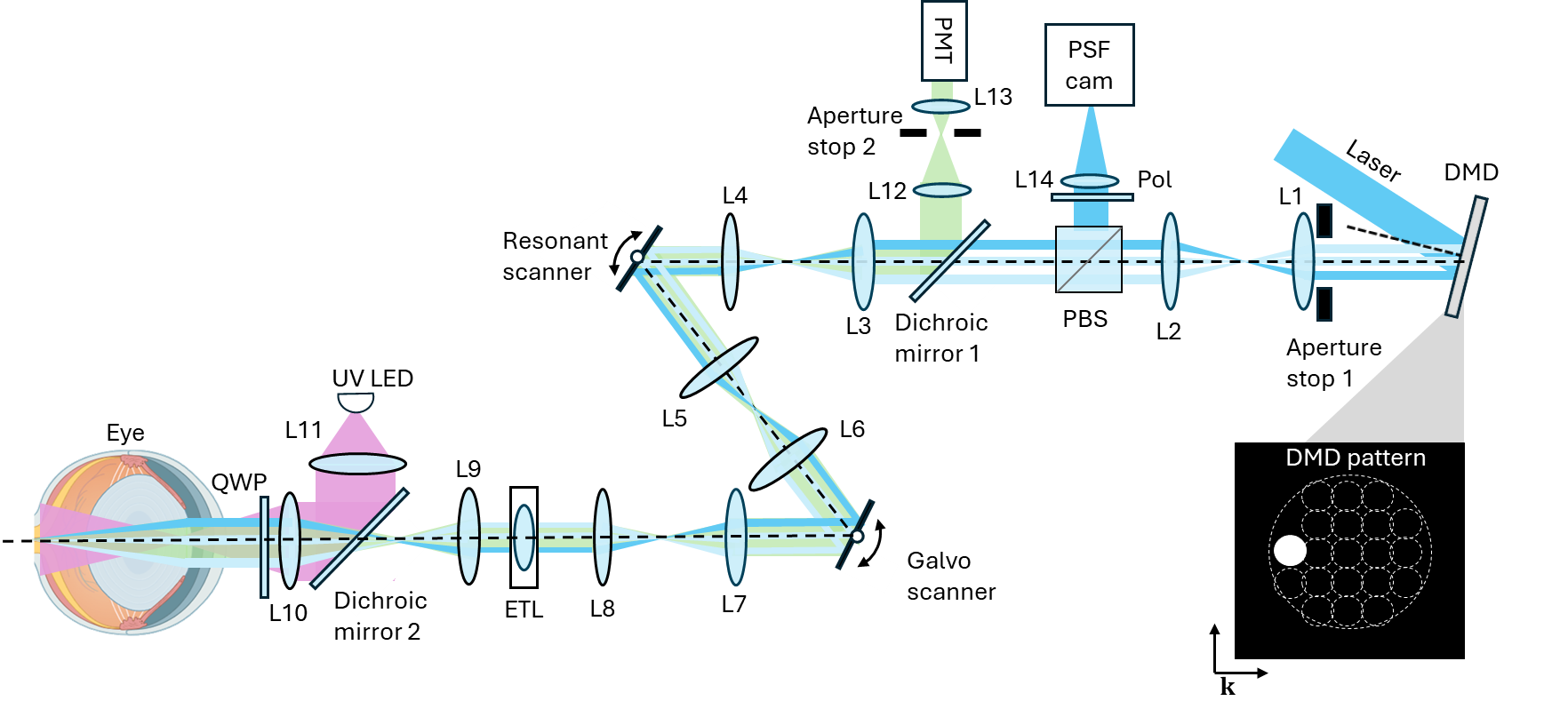

**Figure S1 Schematic of PI-SLO system.**

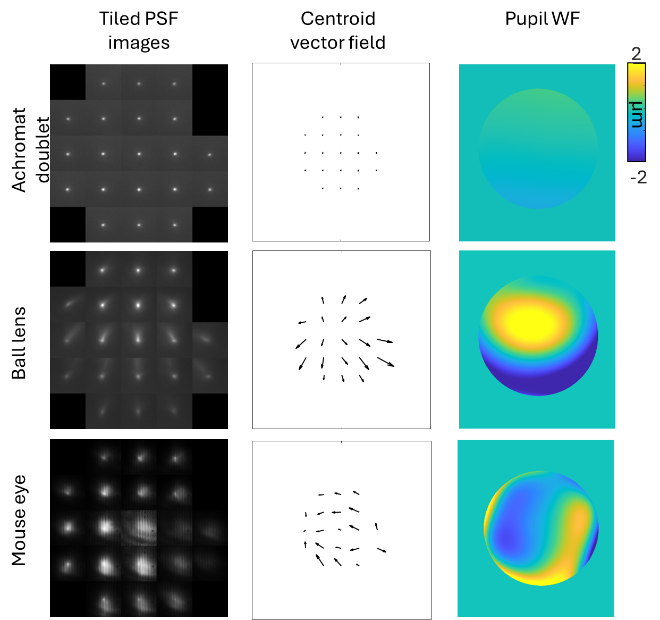

**Figure S2 Synthetic Hartmann-Shack wavefront sensing measurement of an ideal achromatic doublet (Top), a ball lens (Middle), and a living mouse eye (Bottom).**

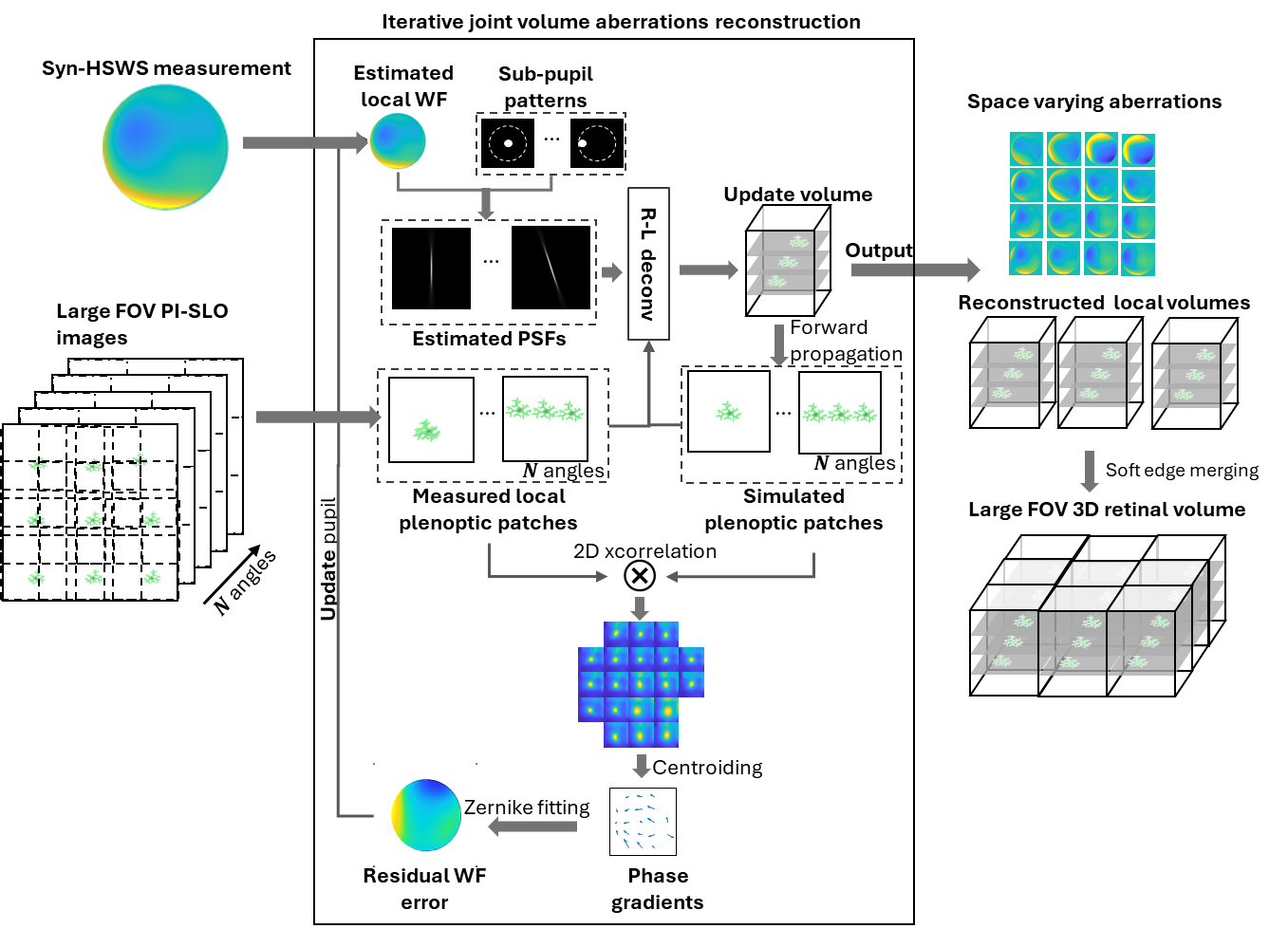

**Figure S3 Illustration of 3D reconstruction routine**

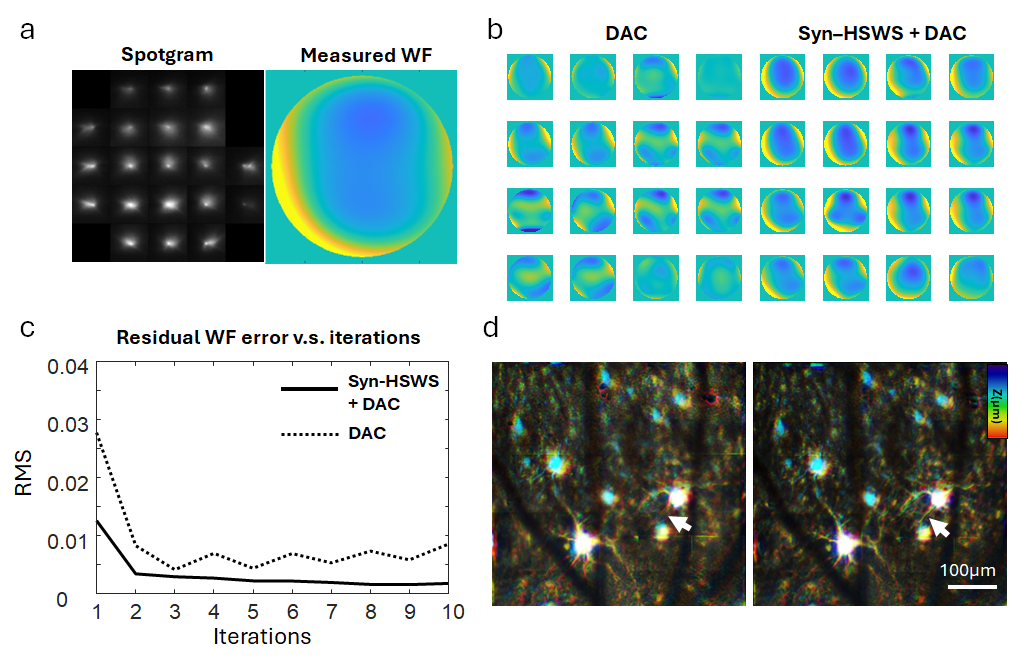

**Figure S4 Sync-HSWF measurement as prior improved reconstruction quality and convergence speed.**

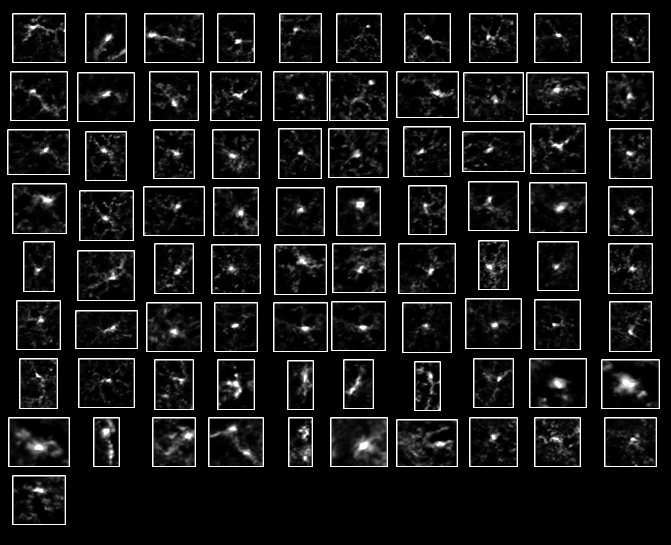

**Figure S5 Manually selected ROIs of 91 single resident immune cells.**

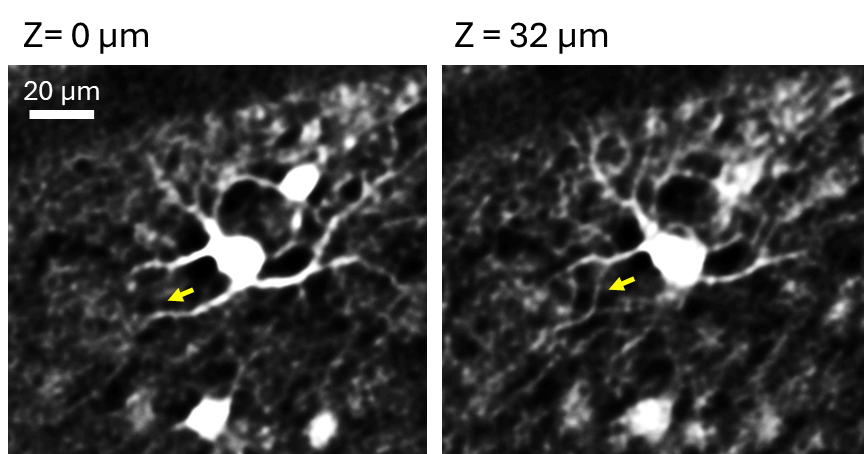

**Figure S6 A bistratified RGC imaged with PI-SLO in a living mouse eye with GCaMP expression.**

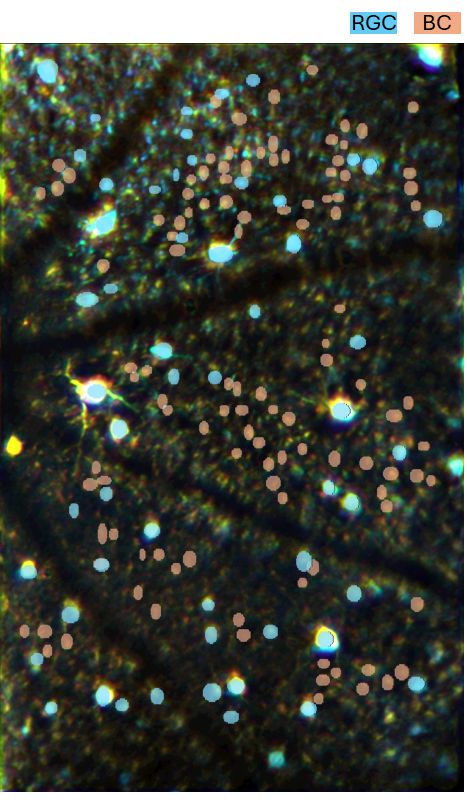

**Figure S7 Manual cell segmentation mask for calcium fluorescent signal extraction**

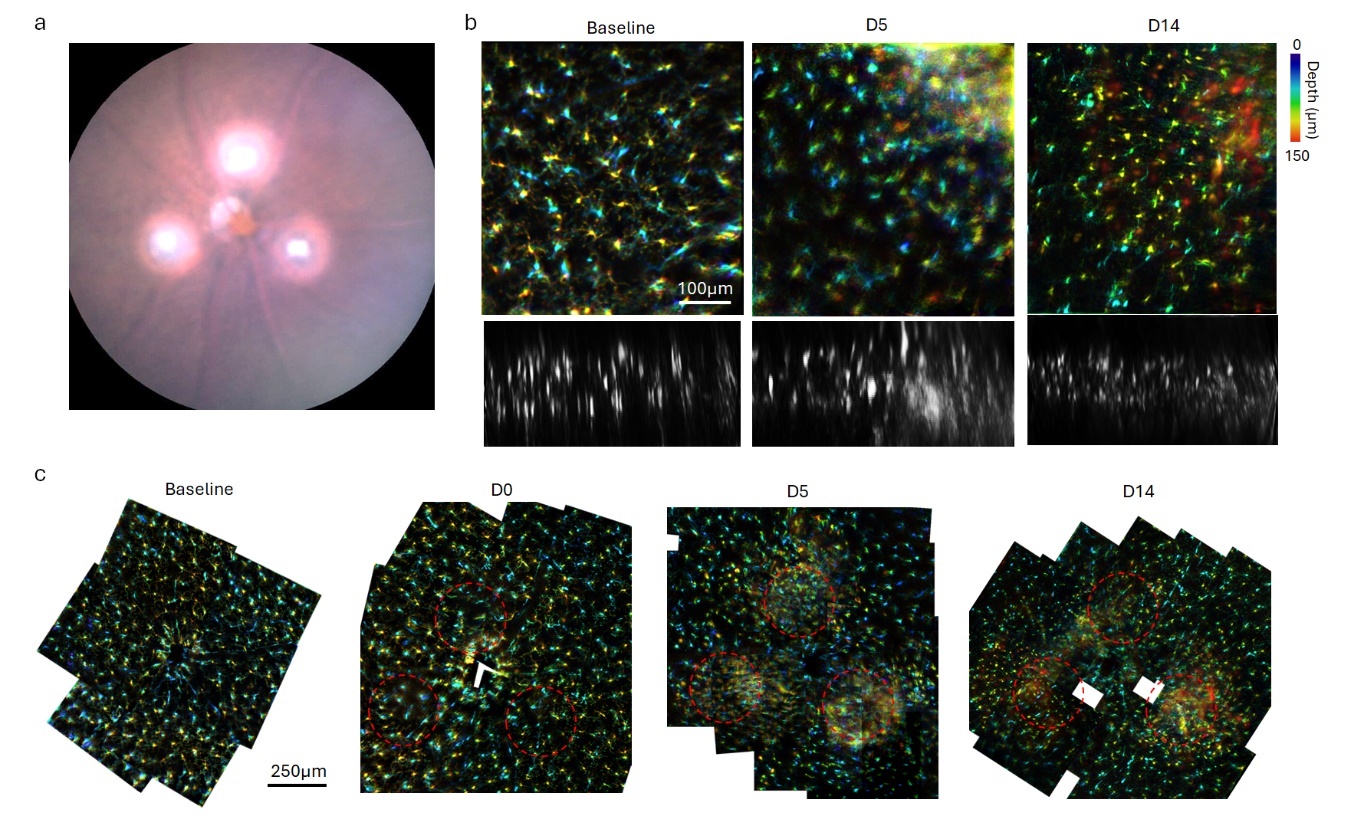

**Figure S8 Laser-induced choroidal neovascularization (CNV) mouse model imaging.**
